# Effects of packetization on communication dynamics in brain networks

**DOI:** 10.1101/2022.06.30.498099

**Authors:** Makoto Fukushima, Kenji Leibnitz

**Affiliations:** Graduate School of Advanced Science and Engineering, Hiroshima University, Hiroshima, Japan; Center for Information and Neural Networks, National Institute of Information and Communications Technology, Osaka, Japan; Graduate School of Information Science and Technology, Osaka University, Osaka, Japan

## Abstract

Computational studies in network neuroscience build models of communication dynamics in the connectome that help us understand the structure-function relationships of the brain. In these models, the dynamics of cortical signal transmission in brain networks are approximated with simple propagation strategies such as random walks and shortest path routing. Furthermore, the signal transmission dynamics in brain networks are associated with the switching architectures of engineered communication systems (e.g., message switching and packet switching). However, it has been unclear how propagation strategies and switching architectures are related in models of brain network communication. Here, we investigate the effects of the difference between packet switching and message switching (i.e., whether signals are packetized or not) on the transmission efficiency of the propagation strategies when simulating signal propagation in a macaque brain network. The results show that packetization decreases the efficiency of the random walk strategy and does not change the efficiency of the shortest path strategy, but increases the efficiency of more plausible strategies for brain networks that balance between communication speed and information cost. This finding suggests an advantage of packet-switched communication in the connectome and provides new insights into modeling the communication dynamics in brain networks.

## Introduction

Methodological advances have enabled mapping the complete set of white matter structural connections (the connectome) in the mammalian brain (Hagmann et al., 2008; Markov et al., 2014; Oh et al., 2014; Stephan et al., 2001). Existing computational studies have investigated the communication dynamics in the connectome by modeling the flow of abstract discrete signals in the network of structural brain connections (Abdelnour, Voss, & Raj, 2014; Crofts & Higham, 2009; Mišić, Sporns, & McIntosh, 2014; see Avena-Koenigsberger, Misic, & Sporns, 2018 for a review). Communication models typically rely on simple approximations of the brain dynamics, while edgewise communication metrics derived from these models explain empirical properties of functional imaging data, such as resting-state functional connectivity weights (Goñi et al., 2014; Mišić et al., 2015).

In communication models for the connectome, the dynamics of signal transmission have been approximated with various propagation strategies. The physiological plausibility of propagation strategies has been discussed in terms of communication speed and information cost (Avena-Koenigsberger et al., 2018; Seguin, van den Heuvel, & Zalesky, 2018). The speed of interregional communication in the cortex needs to be high enough to realize complex brain functions. At the same time, the information cost, i.e., the amount of information required for a cortical signal to propagate toward its destination region (node), should be limited to the extent that no knowledge of the global network topology is used (Avena-Koenigsberger et al., 2018). Several network metrics in connectomics (Rubinov & Sporns, 2010) assume models in which communication takes place through the shortest paths between source and destination nodes; however, this requires centralized knowledge of the global network topology. Alternative models are built on the basis that nervous systems are decentralized. The simplest of such models is the random walk (Mišić, Sporns, & McIntosh, 2014), where a signal randomly propagates from a node to one of its neighboring (structurally connected) nodes. The random walk (RW) and shortest path (SP) strategies are the limiting extremes in terms of communication speed and information cost. With RW, no knowledge is required for propagation, but communication can be slow. With SP, communication is typically fast, but knowledge of the global network topology is necessary. Avena-Koenigsberger et al. (2019) proposed a biased RW strategy that balances between communication speed and information cost.

In parallel with the discussion of propagation strategies, there is an attempt to describe communication in the connectome using an internet metaphor (Graham, 2014, 2021; Graham & Rockmore, 2011). In their model, brain network communication is realized via the propagation of packets split from each signal in a form corresponding to packet switching (Kleinrock, 1976), the switching architecture used in the internet. Physical implementations of packet switching in the brain have not been established; for instance, how individual packets are correctly reassembled at the destination has remained elusive. Nevertheless, the packet-switched communication model has several potential advantages (Graham & Rockmore, 2011), including its ability to reroute network traffic to avoid congestion at hub nodes in, e.g., structural brain networks (Sporns, Honey, & Kötter, 2007) and its efficiency in systems relying on temporally sparse bursts of communication through, e.g., neural spiking activity (Baddeley et al., 1997; Tolhurst, Smyth, & Thompson, 2009). Graham and Rockmore (2011) contrasted packet switching with its precursors in telecommunication systems: message switching and circuit switching. In packet switching and message switching, signals are transmitted in the network toward their destination nodes under a given propagation strategy. The difference between these switching architectures is whether each signal is split into individual packets or not in transit. In circuit switching, a path of connections between source and destination nodes is first established for a signal and then used exclusively to transmit this signal.

While existing models vary in how they approximate the communication dynamics in the connectome regarding propagation strategies and switching architectures, the relationship between these two model components has not been sufficiently explored in the literature. Here, we investigate how different switching architectures affect the performance of propagation strategies in brain network communication models. We exclude circuit switching because its assumption that paths are established before transmission is incompatible with the propagation strategies we consider. Therefore, we focus on examining how splitting signals into packets changes propagation performance. We evaluate performance using transmission efficiency in discrete-event simulations of signal propagation in a macaque brain network. We start with the previous communication model for the connectome in Mišić, Sporns, and McIntosh (2014) for simulating message switching and modify it to simulate packet switching. As propagation strategies, we use RW, SP, and modified versions of RW that have intermediate properties between RW and SP in terms of communication speed and information cost. We demonstrate the effects of packetization on the transmission efficiency of these propagation strategies and discuss how our findings provide new insights into the physiological plausibility of brain network communication models.

## Results

### Simulations of signal propagation in the connectome

We simulated the propagation of discrete signals in the connectome derived from the Collation of Connectivity data on the Macaque brain (CoCoMac) database (Harriger et al., 2012; Kötter, 2004; Modha & Singh, 2010; Stephan et al., 2001) (Figure 1A). We used the same network data of structural brain connectivity as in previous studies (Mišić, Goñi, et al., 2014; Mišić, Sporns, & McIntosh, 2014). These studies described the communication dynamics in the connectome using a discrete-event simulation model with a queueing system (Banks & Carson II, 1984) (Figure 1B). We followed their approach in our simulations of signal propagation with message switching and the RW strategy, where signals (messages) of equal length were randomly generated in time and space at source nodes in the brain network. To each message, a destination node was randomly assigned. When RW was used as the propagation strategy, the message was randomly transmitted to one of the neighboring nodes with the same probability until it arrived at its destination node. When other propagation strategies were used, the message was transmitted to a neighboring node under a given propagation strategy toward its destination node. Following the previous studies of Mišić and colleagues, nodes in the network were modeled as servers with finite buffer capacity (maximum number of slots *H* = 20; see the Supplementary Results section for the results with no buffer size limit). Once a message arrived at a node, it started service if there was no other message occupying that node, or it was stored in the buffer otherwise, forming a queue (see Figure 1B, bottom). When a previously occupying message finished its service, the newest message was to start its service (last-in-first-out queueing; Kleinrock, 1976; Banks & Carson II, 1984). A message was removed from the network when it reached its destination node or when it was the oldest message in a fully occupied buffer at which another message newly arrived. We used the same arrival rate *λ* (average number of messages generated per time unit) and service rate *µ* (inverse of average service time) as in the previous studies of Mišić and colleagues (*λ* = 0.01 and *µ* = 0.02). Since the ratio *λ*/*µ* governed the system dynamics, we also report the results with decreased and increased *λ* with fixed *µ* in the Supplementary Results section.

**Figure 1.**
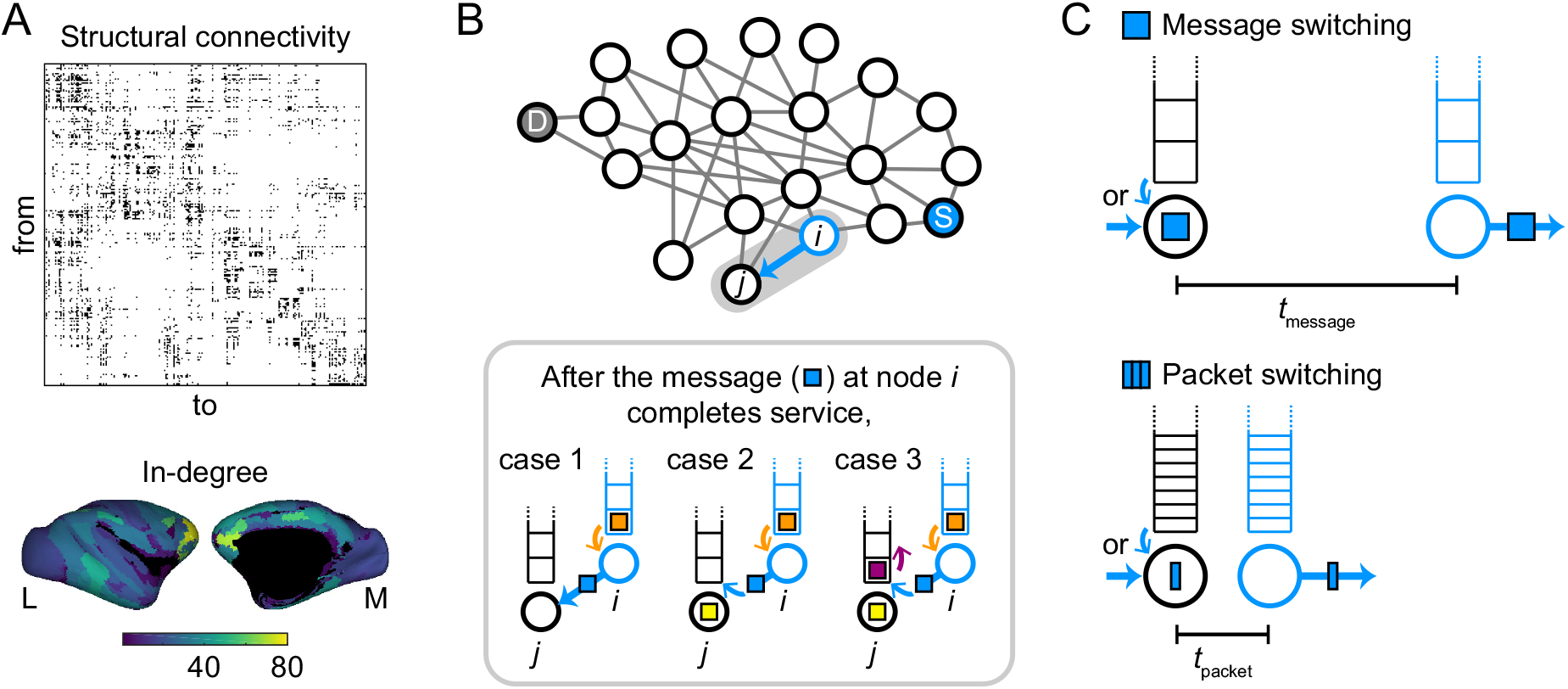
Simulation models of signal propagation in the connectome. (A) Adjacency matrix of the network of structural connectivity (242 cortical nodes, 4, 090 binary and directed edges) used in simulations (top) and rendering of its in-degree (the number of incoming connections to a node) on cortical surfaces (bottom left, lateral view; bottom right, medial view). The surface maps only show the in-degrees of the 187 nodes for which spatial coordinates are available (Harriger et al., 2012). (B) Schematic description of the discrete-event simulation. In simulations of signal propagation with message switching (Mišić, Goñi, et al., 2014; Mišić, Sporns, & McIntosh, 2014), every signal (message) was associated with a randomly generated pair of source node S and destination node D. Once the message (colored light blue) completes service at node *i*, it is transmitted to node *j* and the message (orange) in the buffer of node *i* starts service at node *i*. If there is no signal at node *j* (case 1), the light blue message immediately starts service at node *j*. If another message (yellow) is in service at node *j* (cases 2 and 3), the arriving light blue message is stored in the front of the buffer of node *j*. In case 3, where a message (purple) is in the front of the buffer of node *j*, the light blue message is stored in the front and the purple message is moved back. (C) Modification of the discrete-event simulation model to simulate signal propagation with packet switching. When a message is split into *n* equally sized packets, the service rate and the maximum number of slots in a buffer for a packet are specified as *n* times as large as those for a message. In this case, the service time for a packet (*t*_packet_) is on average that for a message (*t*_message_) divided by *n*.

In the discrete-event simulation model of Mišić and colleagues, an entire signal (message) was trans-mitted to a neighboring node at once by message switching. We modified this model to simulate signal propagation in the connectome with packet switching (Figure 1C). In the modified model, a message was divided into *n* equally sized packets. We ran simulations with *n* = 5 as the default for all packets and then *n* = 3 and *n* = 10 to check the reproducibility of the results (see the Supplementary Results section). Packets of the same message were simultaneously generated at the source node, and each packet was independently transmitted to a neighboring node under a given propagation strategy. All packets of the same message had the same destination but could take different routes to reach it. To take the size differences of messages and packets into account, we scaled the service rate *µ* and the maximum number of slots in the buffer *H* by *n* (see Figure 1C). Further details of the simulations are described in the Materials and Methods section.

### Effects of packetization on transmission efficiency

After simulating signal propagation in the connectome, we compared the transmission efficiency of each propagation strategy between message switching and packet switching. Transmission efficiency was evaluated by measuring the elapsed time to transmit 100 messages or their corresponding packet sets between source and destination nodes, i.e., the duration from when the first message or packet was generated in the network until the transmission of 100 messages or packet sets to their destinations was completed. A shorter elapsed time corresponds to a higher transmission efficiency.

We first evaluated the transmission efficiency of RW, SP, and informed RW (iRW) that uses local information only from neighboring nodes for propagation (Figure 2A). We implemented three versions of iRW. One is a version with a rule of avoiding busy neighboring nodes in transit (iRW_a_). In this version, a message or packet is randomly transmitted to a neighboring node with no message or packet in service or, if such a node does not exist, to a neighboring node with the minimum queue length in the buffer. Another version is with a rule of deterministic transmission to the destination node if it is one of the neighboring nodes (iRW_d_). The third version is with the combination of the above two rules (iRW_a+d_).

**Figure 2.**
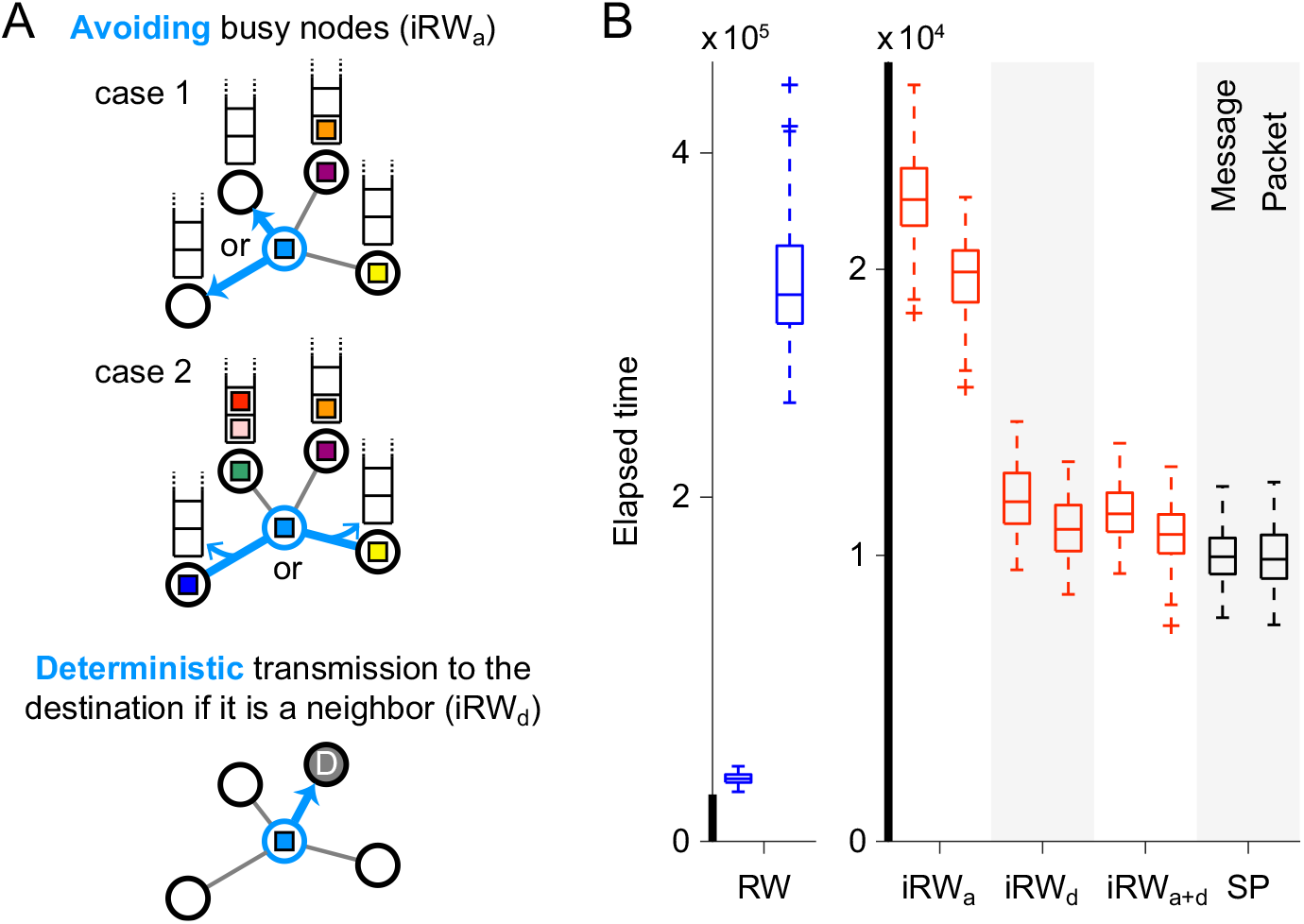
Description of iRW and results of elapsed time comparison for RW, iRW, and SP. (A) Schematic description of iRW. In iRW with the rule of avoiding busy nodes (iRW_a_ and iRW_a+d_), a message or packet (colored in light blue) is transmitted to a neighboring unoccupied node if it exists (case 1) or to the neighboring node with the minimum queue length in the buffer (case 2). In iRW with the rule of deterministic transmission to the neighboring destination node (iRW_d_ and iRW_a+d_), a message or packet is deterministically transmitted to the destination node if it is included in the set of neighboring nodes. (B) Elapsed time for transmitting 100 messages or packet sets in the connectome with RW, iRW_a_, iRW_d_, iRW_a+d_, and SP. The left and right columns for each propagation strategy in the boxplots show the elapsed times of message switching and packet switching, respectively (we ran 100 simulations to obtain the distribution of the elapsed time in each column). Pairs of columns for each propagation strategy are shown in red (blue) when the elapsed time of packet switching was shorter (longer) than that of message switching or in black when no significant differences were observed between them (*p <* 0.05, FDR corrected). The thick black lines in the vertical axes indicate the same range of the elapsed time.

Figure 2B shows the elapsed time for transmitting 100 messages or packet sets in the connectome with RW, iRW_a_, iRW_d_, iRW_a+d_, and SP. The elapsed time of packet switching was longer than that of message switching under RW (Mann-Whitney *U* test; *p* = 1.28 *×* 10*^−^*^33^, two-sided, false discovery rate (FDR) corrected by the Benjamini-Hochberg method; Cliff’s delta = 1) and the elapsed times of both switching architectures were similar under SP (*p* = 0.848, FDR corrected, Cliff’s delta = *−*0.016). By contrast, the elapsed time of packet switching was shorter under iRW (iRW_a_, *p* = 1.06 *×* 10*^−^*^24^, FDR corrected, Cliff’s delta = *−*0.847; iRW_d_, *p* = 3.49 *×* 10*^−^*^7^, FDR corrected, Cliff’s delta = *−*0.425; iRW_a+d_, *p* = 9.80 *×* 10*^−^*^7^, FDR corrected, Cliff’s delta = *−*0.404), indicating that splitting a message into packets improved the transmission efficiency of iRW. All versions of iRW only need local information from neighboring nodes, but their elapsed times (especially those with iRW_d_ and iRW_a+d_) were much closer to the elapsed time with SP than that with RW. These results suggest that packetizing signals in the connectome further degrades the transmission efficiency of slow strategies (e.g., RW), but conversely improves the efficiency of strategies that balance between communication speed and information cost (e.g., iRW).

We next evaluated the transmission efficiency of biased RW (bRW) (Avena-Koenigsberger et al., 2019) across its control parameter *c* that can shift the propagation behavior of bRW between RW and SP. Figure 3A illustrates the transition probabilities under bRW in a toy example case. When a message or packet is transmitted to one of the neighboring nodes whose path lengths to the destination node are (i) one, (ii) two, and (iii) three (Figure 3A, top), respectively, the transition probability to (i) is near 1/3 (one over the number of neighboring nodes; RW-like behavior) when *c* = 0.01 and near 1 (SP-like behavior) when *c* = 10 (Figure 3A, bottom). For the formal definition of the transition probability under bRW, see the Materials and Methods section. The shift of propagation behavior over the spectrum of *c* can be associated with the change in the level of availability of global information about the network structure (Avena-Koenigsberger et al., 2019). Therefore, although the shortest path lengths are required to compute the transition probabilities in the implementation, we regarded bRW with an intermediate to low range of *c* as an effective propagation strategy with respect to the information cost.

**Figure 3.**
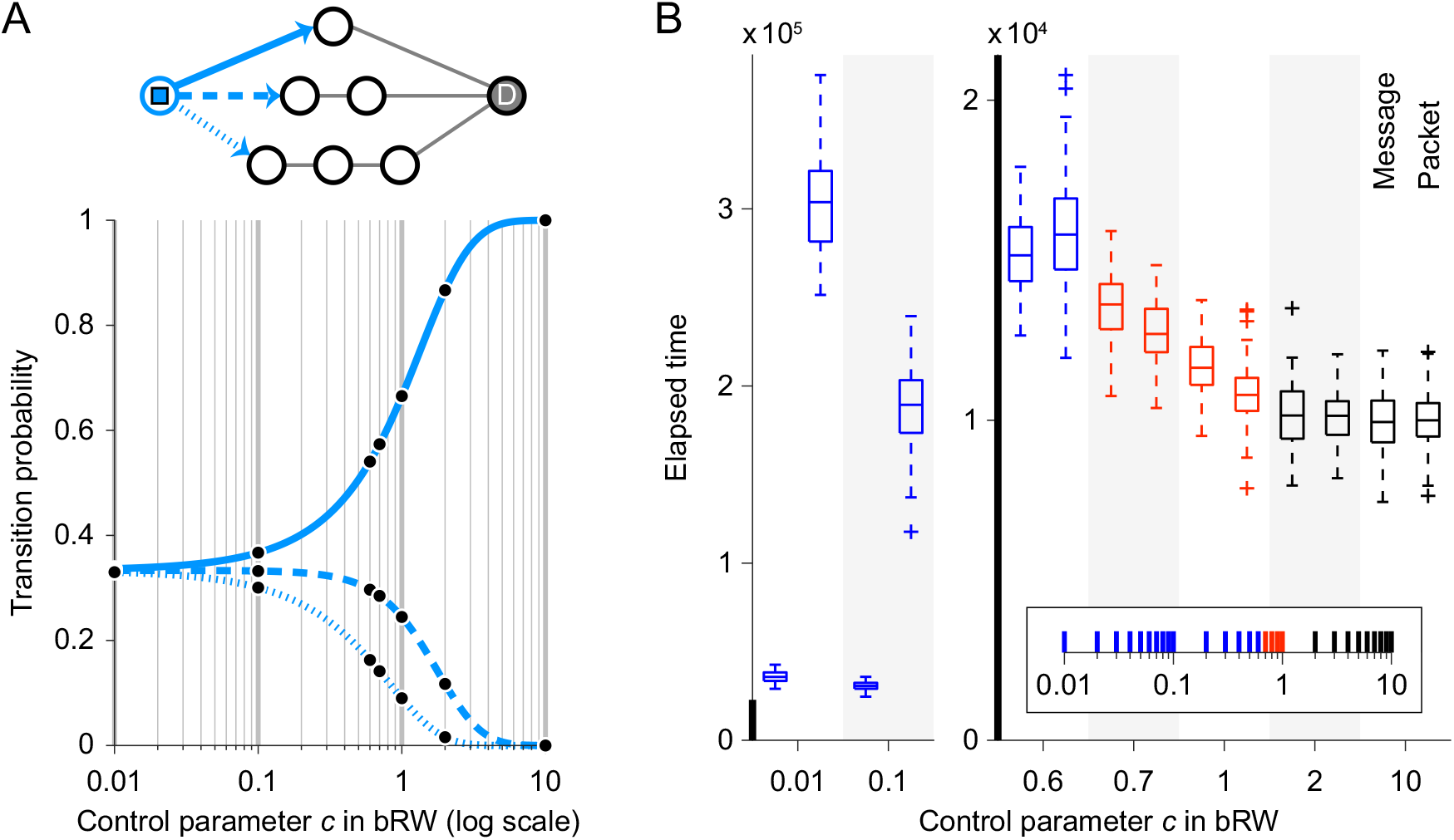
Description of bRW and results of elapsed time comparison for bRW. (A) Transition probability to neighboring nodes in a toy example case under bRW over the spectrum of the control parameter *c*. The transition probabilities are shown for the message or packet at the leftmost node (in the network shown above the plot) that has three neighboring nodes whose path lengths to the destination node D are one (top), two (middle), and three (bottom). The line types of the arrows toward the neighboring nodes correspond to the line types in the plot of the transition probabilities. The black dots in the plot indicate the transition probabilities at *c* for which the boxplots of the elapsed time are shown in panel B. (B) Elapsed time for transmitting 100 messages or packet sets in the connectome with bRW over the spectrum of *c*. The boxplots are colored in the same manner as in Figure 2B (blue: longer elapsed time of packet switching; red: shorter elapsed time of packet switching; black: no time difference between message switching and packet switching) (*p <* 0.05, FDR corrected for the 28 comparisons at *c ∈ {*0.01, 0.02, …, 0.09, 0.1, 0.2, …, 0.9, 1, 2, …, 9, 10*}*). The inset located at the bottom right summarizes the statistical results at all the considered values of *c*.

In Figure 3B, we show the elapsed time for transmitting 100 messages or packet sets in the connectome under bRW across the spectrum of *c*. When *c* was relatively small, the elapsed time of packet switching was longer than that of message switching (e.g., *c* = 0.01, *p* = 5.98 *×* 10*^−^*^34^, FDR corrected, Cliff’s delta = 1) as seen in the case with RW. When *c* was relatively large on the other hand, no significant difference was observed between the elapsed times of message switching and packet switching (e.g., *c* = 10, *p* = 0.923, FDR corrected, Cliff’s delta = *−*0.014) as in the case with SP. The elapsed time of packet switching was shorter than that of message switching when *c* was specified within an intermediate range (e.g., *c* = 0.7, *p* = 2.35 *×* 10*^−^*^8^, FDR corrected, Cliff’s delta = *−*0.463; *c* = 1, *p* = 1.26 *×* 10*^−^*^9^, FDR corrected, Cliff’s delta = *−*0.504). In this range, the speed of communication was high and comparable to the speed under SP. These results also suggest that packetization in the connectome improves the transmission efficiency of speed- and cost-balanced propagation strategies.

We confirmed that the results of the elapsed time comparisons were robust against various changes applied to the default simulation setting. We describe these changes and show the results obtained with them in the Supplementary Results section and Figures S1–S6.

### Communication metrics in simulations

So far, we have seen that packetization has different effects on the transmission efficiency of RW, iRW/bRW with an intermediate range of *c*, and SP. In this section, we investigate its reason by computing nodewise communication metrics derived from the simulated signal propagation in the connectome. We employed communication metrics computed in the previous studies (Mišić, Goñi, et al., 2014; Mišić, Sporns, & McIn-tosh, 2014): the proportion of time that a node was busy (*utilization*) and the mean number of signals (messages or packets) at a node (*node contents*) (see the Materials and Methods section for their formal definitions). To properly compare the node contents between message switching and packet switching, we divided the value of the node contents at a node by its total capacity (= the maximum buffer size plus one) for normalization.

Figure 4 presents the relations between the computed communication metrics and the in-degree (the number of incoming connections) of the structural brain network used in the simulations (see Figure 1A). In SP and bRW with *c* = 2, utilization was almost identical between message switching and packet switching, which would explain why the elapsed times of both switching architectures were essentially the same under these propagation strategies. In iRW and bRW with *c* = 0.7 and 1, utilization was greater and normalized node contents were smaller for packet switching in most nodes, indicating that packets propagated more diversely across the network compared to messages. This result suggests that packetization allows a more efficient use of the entire network, which reduces the elapsed time with these propagation strategies. In RW and bRW with *c* = 0.1 and 0.6, normalized node contents for packet switching were more than 1.5 times larger than those for message switching in a few high in-degree nodes indicated by arrows and ellipses in Figure 4B. These hub nodes that gathered many packets were macaque brain areas 23c (RW and bRW with *c* = 0.1) in the cingulate cortex, 12o (RW), 13a (RW), 32 (RW and bRW with *c* = 0.1), and 46 (RW and bRW with *c* = 0.1 and 0.6) in the prefrontal cortex (see Figure 4c for the positions of these areas on the cortical surface). The utilization metric averaged over these nodes for packet switching was 0.98, which means that the nodes were almost always occupied by packets. These results indicate that a few hub nodes, which played the role of bottlenecks in the connectome, were responsible for the lower efficiency of packet-switched communication under the slow propagation strategies.

**Figure 4.**
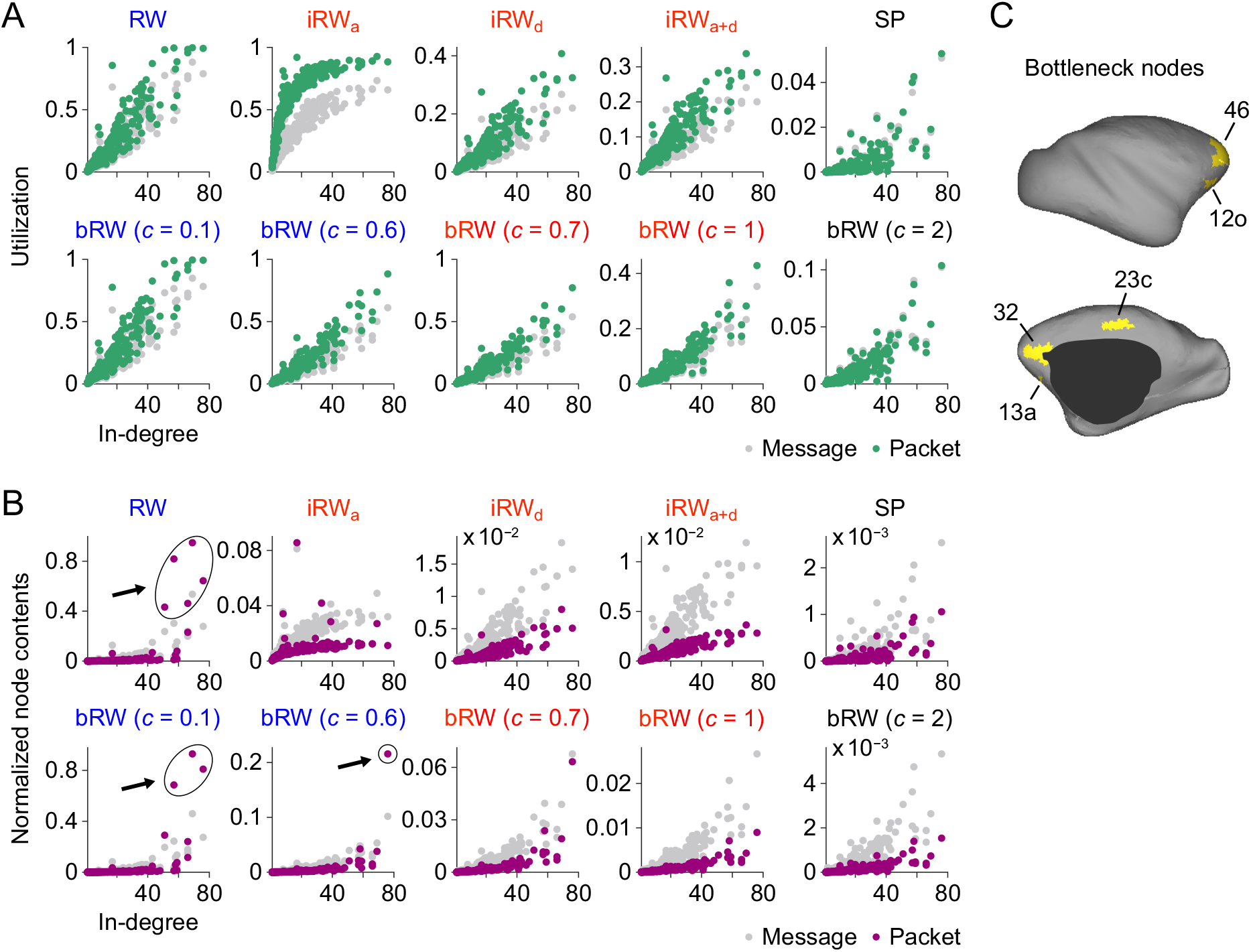
Nodewise communication metrics in the simulations of signal propagation. (A) Relation between the utilization metric and the in-degree of the structural brain network (each dot represents a node). Utilization was averaged over 100 simulation samples. Scatter plots for message switching (gray) and packet switching (green) are presented for each propagation strategy. The text color of each strategy corresponds to the color of the boxplots in Figures 2B and 3B. (B) Relation between the normalized node contents metric and the in-degree of the structural brain network. Normalized node contents were averaged over 100 simulation samples. Arrows and ellipses in the scatter plots indicate bottleneck nodes whose normalized node contents for packet switching (purple) were more than 1.5 times larger than those for message switching (gray). (C) Spatial positions of the bottleneck nodes on the cortical surface of the macaque brain.

### Simulations using surrogate networks

Finally, we investigate how the network properties of the connectome affect the results of the elapsed time comparisons for message switching and packet switching. For this purpose, we simulated signal propagation in surrogate networks constructed by rewiring edges in the original structural brain network. We employed surrogate networks in which locations of edges were fully randomized (Figure 5A; no hub nodes existed) and those in which random edge rewiring was constrained to preserve the in-degree and out-degree (the numbers of incoming and outgoing connections) of nodes in the original network (Figure 5B; hub nodes remained).

**Figure 5.**
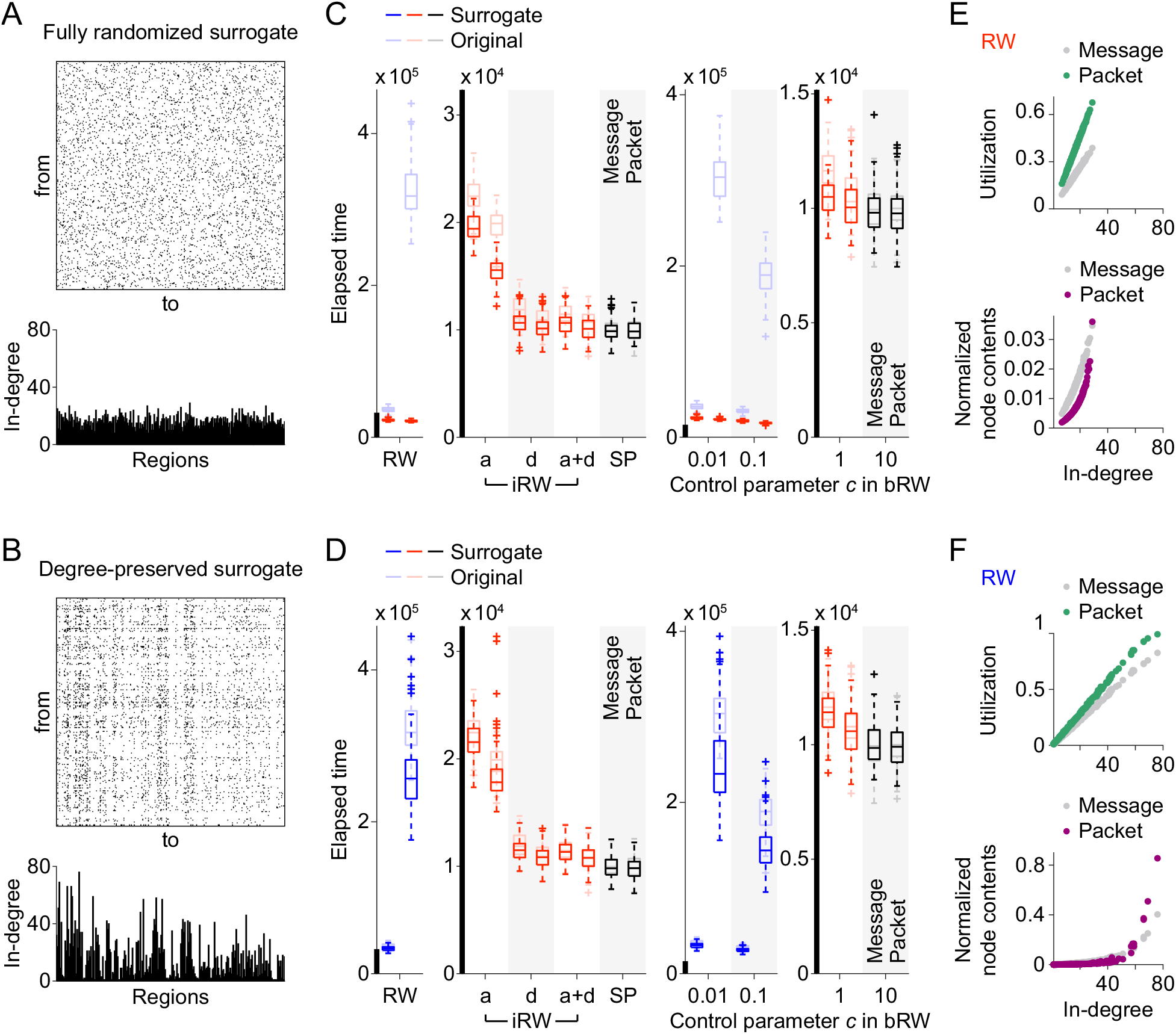
Results of the simulations using surrogate networks. (A) Adjacency matrix (top) and in-degree (bottom) of a fully randomized surrogate network. (B) Adjacency matrix (top) and in-degree (bottom) of a degree-preserved surrogate network. (C) Elapsed time for transmitting 100 messages or packet sets in the simulations using the fully randomized surrogate networks. (D) Elapsed time for transmitting 100 messages or packet sets in the simulations using the degree-preserved surrogate networks. The elapsed times obtained from the original brain network are also shown for reference in panels C and D. (E) Relation between utilization (top) or normalized node contents (bottom) and in-degree, derived from 100 samples of the fully randomized surrogate networks. (F) Relation between utilization (top) or normalized node contents (bottom) and in-degree, derived from 100 samples of the degree-preserved surrogate networks. Utilization and normalized node contents were averaged over all samples. When averaging these metrics, nodes were sorted in an ascending order of their in-degrees.

Figure 5C shows the elapsed time for transmitting 100 messages or packet sets in the simulations using the fully randomized surrogate networks. In contrast to the results obtained from the original brain network, packetization shortened the elapsed time under RW and bRW with *c* = 0.1 and 0.01 (Figure 5C; for the boxplots zoomed in along the vertical axis, see Figure S7). Nodes acting as bottlenecks in packet switching disappeared, where no node exhibited a utilization close to one (Figure 5E, top) or normalized node contents for packet switching more than 1.5 times larger than those of message switching (Figure 5E, bottom). On the other hand, with the degree-preserved surrogate networks, the elapsed time of packet switching was longer than that of message switching under the same strategies (Figure 5D) and the bottleneck nodes also remained (Figure 5F). However, the effects of packetization on the elapsed time were less pronounced compared to the case in the simulations using the original brain network (Figure 5D). These results suggest that hub nodes in particular, but also the entire network topology of the connectome, contribute to slowing down RW-style communication through packetization.

## Discussion

While propagation strategies and switching architectures are key components of brain network communication models, their relationships among each other have been unclear. To address this issue, we investigated how the difference between packet switching and message switching (i.e., whether signals are packetized or not) affects the performance of propagation strategies in the connectome. By performing simulations of signal propagation in a macaque brain network, we found that packetization in the connectome degraded the transmission efficiency of slow propagation strategies such as RW and did not change the efficiency of costly SP, but improved the efficiency of strategies that balanced between communication speed and information cost. Nodewise communication metrics in the simulations indicate that packetization caused some high-load bottleneck nodes to appear in the connectome under the slow strategies but reduced the load on most nodes under the balanced strategies by allowing more nodes to process packetized signals. Surrogate network analysis suggests that the degraded efficiency through packetization in the slow strategies was not only due to the presence of high-degree hub nodes, but also due to the network topology of the connectome in the brain.

The findings in this study provide new insights into the physiological plausibility of the two major components of brain network communication models: propagation strategies and switching architectures. RW and SP are classic propagation strategies often assumed in existing models and metrics for brain network analysis (Mišić, Sporns, & McIntosh, 2014; Rubinov & Sporns, 2010). However, due to the slow communication speed of RW and the high information cost of SP, propagation strategies that balance the speed and cost have been considered more appropriate for use in brain network communication models (Avena-Koenigsberger et al., 2018; Seguin et al., 2018). Our finding that packetization degrades signal transmission efficiency of RW in the connectome further supports this view. Moreover, we have shown that packet switching with balanced strategies is more advantageous than message switching for achieving efficient communication in the connectome. Message switching and packet switching are both compatible with the concept of dynamic routing in the brain (Gerraty et al., 2018; Nádasdy, Hirase, Czurkó, Csicsvari, & Buzsáki, 1999; Palmigiano, Geisel, Wolf, & Battaglia, 2017). While message switching has been implemented explicitly or implicitly in many models of brain network communication (e.g., Mišić, Sporns, & McIntosh, 2014), Graham and Rockmore (2011) introduced packet switching to network neuroscience research and theoretically assessed its potential advantages (e.g., rerouting ability) over message switching for interregional communication in the brain. The present study has revealed the merit of packet switching based on empirical simulations of signal propagation in the connectome. Although there are challenges in explaining how packet-switched communication schemes are implemented in the nervous system (as noted below regarding the limitations of modeling based on packet switching), the increase in transmission efficiency of balanced propagation strategies through packetization strengthens the value of packet switching as an element of communication models for brain networks.

A limitation of packet switching as a component of brain network communication models is that it is difficult to explain how packets in the same message are reassembled at their destination node in the connectome. Since these packets can take different paths from their source node (Graham & Rockmore, 2011), the order in which the packets arrive at the destination can be reversed. To address this problem, an additional modeling assumption is necessary, namely that the original message is split into packets that can be reassembled in any order. One example following this assumption would be to packetize a message into different features rather than temporal fragments. An alternative solution is to model packet switching that does not allow packet overtaking. Although it reduces the flexibility of propagation, no packet overtaking can be implemented by, for example, introducing the assumption that all packets in the same message follow the path of their forerunners and replacing the last-in-first-out queueing system with the first-in-first-out one (Banks & Carson II, 1984; Kleinrock, 1976). Even with such a model of packet switching with no packet overtaking, we confirmed that packetization improved the transmission efficiency of speed- and cost-balanced propagation strategies (see Figure S6). This alternative model could alleviate concerns about explaining the physical implementation of packet reassembly in brain networks.

Another problem with the packet-switched communication model is that packetization can reduce the reliability of communication in brain networks. Errors may occur when packets are reassembled at the destination node, although we did not include such errors in the simulations. In addition, when signals are congested in the connectome, packetizing signals may make it more difficult to complete the transmission of the entire portion of individual signals to their destinations, since the possibility of a packet being lost along the way would be greater than that of a message due to the increased number of discrete instances propagating through the network. Our simulations confirmed that slow propagation strategies such as RW caused excessive concentration of packets at bottleneck nodes, resulting in greater chances of information loss in the buffers at these nodes. The buffers that reflect the function of working memory (Funahashi, 2015; Goldman-Rakic, 1996) contribute to reliable communication, but additional mechanisms may be needed to promote the reliability of packet-switched communication. One solution is to introduce redundancy into brain network communication (Avena-Koenigsberger et al., 2017; Bettinardi et al., 2017). Duplicating signals and transmitting them through multiple paths in the connectome help improve the reliability of communication when signals collide destructively at nodes in brain networks (Hao & Graham, 2020). Duplication could be introduced into packet switching by, for example, copying several important packets to increase the chance of signal recovery at the destination node. Such a redundant propagation scheme would be necessary to make packet-switched communication more reliable even in the presence of signal congestion in the connectome.

There are also other methodological limitations to this study. First, our simulations are based on an abstract modeling approach to cortical signal propagation that is typical of many previous computational studies of brain network communication (Abdelnour et al., 2014; Avena-Koenigsberger et al., 2018, 2019; Crofts & Higham, 2009; Goñi et al., 2014; Mišić et al., 2015; Mišić, Goñi, et al., 2014; Mišić, Sporns, & McIntosh, 2014; Seguin et al., 2018). We focused on this approach because it allows us to trace the propagation of individual signaling units in the connectome. However, the results should be interpreted with the caveat that the models used here are simpler than the more realistic biophysical models of neuronal population dynamics that can reproduce ongoing large-scale brain activities (Breakspear, 2017; Cabral, Hugues, Sporns, & Deco, 2011; Deco et al., 2017; Deco & Jirsa, 2012; Fukushima & Sporns, 2018, 2020; Honey, Kötter, Breakspear, & Sporns, 2007; Honey et al., 2009; Pope, Fukushima, Betzel, & Sporns, 2021). In addition, all nodes and edges were treated equally in the models, despite the fact that brain regions vary in size and their structural connections vary in length and width. The models were configured in this way to avoid making even stronger assumptions about how these variations affect the dynamics of communication in brain networks. Second, only a limited number of propagation strategies were used in the simulations. A single propagation strategy was assumed for all communication processes in an individual simulation sample, whereas it is possible that neural systems may combine aspects of multiple propagation strategies (Avena-Koenigsberger et al., 2018; Betzel, Faskowitz, Mišić, Sporns, & Seguin, 2022). Navigation (Seguin et al., 2018) and broadcasting (Mišić et al., 2015) strategies were also not investigated because navigation requires the location information of nodes, which is missing for several nodes in the brain network used in this study, and broadcasting multiplies signals over time, making it difficult to compare results with those obtained from conventional propagation strategies (e.g., RW and SP). Third, the connectome dataset

(Harriger et al., 2012) used in this study is somewhat old. This dataset was chosen because it was used in the previous simulation studies (Mišić, Goñi, et al., 2014; Mišić, Sporns, & McIntosh, 2014), which allowed us to use the same parameter values when simulating message-switched communication in the connectome. Future work could run the simulations on more recent connectome data, such as those developed by Oh et al. (2014) for the mouse brain and/or those of Chen, Zhang, He, and Zhou (2020), who combined the connectivity information from the CoCoMac database and from Markov et al. (2014) for the macaque brain.

The current study is a first step toward a comprehensive assessment of the physiological plausibility of propagation strategies and switching architectures in brain network communication models. We have demonstrated the advantage of packet switching for communication in brain networks under speed- and cost-balanced propagation strategies; however, this finding only indirectly supports the plausibility of this switching architecture. For more direct evidence, it would be necessary to include evaluations based on empirical functional data (Seguin, Tian, & Zalesky, 2020). A promising approach for such evaluation is to compute the similarity between edgewise communication metrics quantifying the frequency of simulated signal transmission (Mišić, Sporns, & McIntosh, 2014) and the weights of empirical resting-state functional connectivity (Damoiseaux et al., 2006; Fox et al., 2005; Yeo et al., 2011). Computing this similarity, which quantifies the extent to which a communication model can explain empirical functional interactions in the brain, may allow us to evaluate the plausibility of a given combination of propagation strategy and switching architecture. We will conduct this line of future research using the Human Connectome Project dataset (Van Essen et al., 2013), from which we can construct structural and functional brain networks and relate simulated edgewise communication metrics to empirical functional connectivity.

## Materials and Methods

### Connectome data

The connectome data were derived from the CoCoMac database (Kötter, 2004; Stephan et al., 2001) that provides white matter structural connectivity information of the macaque brain reported in existing tract tracing studies. Connectivity information was initially collected by querying this database in Modha and Singh (2010), later refined to obtain a fully connected brain network (Harriger et al., 2012), and used in simulations of signal propagation in Mišić, Sporns, and McIntosh (2014) and Mišić, Goñi, et al. (2014). This structural brain network was comprised of 242 cortical regions (nodes) and 4, 090 directed connections (edges) represented in binary format (connection present = 1 and absent = 0) and contained no self-connections (see Figure 1A).

### Discrete-event simulation and switching architectures

Signal propagation in the connectome was simulated using discrete-event simulation techniques (Banks & Carson II, 1984). In the simulations with message switching, individual signals (messages) were randomly generated in the brain network as a Poisson process with exponentially distributed inter-arrival times (arrival rate *λ* = 0.01 in the default setting) as in Mišić, Sporns, and McIntosh (2014) and Mišić, Goñi, et al. (2014). Poisson arrivals represent statistical fluctuations of the sensory environment (Barlow, 1956; McGill, 1967). To each message, a pair of source and destination nodes were randomly assigned. A message was transmitted to one of the neighboring nodes under a given propagation strategy until it reached its destination node. After reaching the destination node, the message was removed from the network. The time that a message spent at each node (service time) was exponentially distributed (service rate *µ* = 0.02) as in the previous studies of Mišić and colleagues. The ratio of the arrival rate to the service rate (*λ*/*µ*) governed the dynamics of the whole system and was specified so that the number of messages in the network did not monotonically increase during the simulations. If a message arrived at a node that was already occupied by another message, the arrived message was placed in a buffer and formed a queue. Queueing was used to ensure that the dynamics of the messages were interdependent (Liu, 1996). Messages entered the node on a last-in-first-out basis (Banks & Carson II, 1984; Kleinrock, 1976) and a maximum buffer size was imposed (*H* = 20) in the default setting as in the previous studies of Mišić and colleagues. In this case, a message arriving at a fully occupied buffer caused the oldest message in the queue to be ejected and removed from the network. A finite buffer size was used to model imperfect signal transmission (Faisal, Selen, & Wolpert, 2008). The state of the system was updated at non-uniform intervals because of the presence of stochastic variables in inter-arrival times and service times. Figure 1B shows a schematic description of the discrete-event simulation with message switching.

In the simulations with packet switching, packets split from the entire message were transmitted across the network. A message was divided into *n* = 5 equally sized packets in the default setting. A set of *n* packets was generated at a source node and transmitted to neighboring nodes under a given propagation strategy toward their common destination node. Packets in the same message individually propagated over the network in the default setting, such that they could take different routes and arrive at the destination node with different propagation delays. The service rate and the maximum number of slots in a buffer for a packet were specified as *nµ* and *nH*, respectively (Figure 1C).

### Comparison of transmission efficiency

The transmission efficiency of each propagation strategy was compared between message switching and packet switching to investigate the effects of packetization. We evaluated the transmission efficiency by measuring the elapsed time in simulations to transmit 100 messages or their corresponding packet sets to their respective destination nodes in the connectome (the shorter the elapsed time is, the higher the transmission efficiency is). The elapsed time of message switching was defined as the duration from when the first message was generated at a source node until when 100 messages in total arrived at their destination nodes. The elapsed time of packet switching was the duration from when the first set of packets was generated at a source node until when all *n* packets in 100 packet sets arrived at their destination nodes. We ran 100 simulations each for message switching and packet switching. Using these simulation samples, we assessed whether the elapsed time of packet switching was shorter or longer than that of message switching for each of the propagation strategies described below.

### Propagation strategies

We compared the transmission efficiency between message switching and packet switching under the following strategies of signal propagation in the connectome: random walk (RW), shortest path (SP), and informed and biased versions of RW (iRW and bRW) that have intermediate properties between RW and SP in terms of communication speed and information cost.

In RW, a message or packet was randomly transmitted to one of the neighboring nodes of the current node with an equal probability. In this strategy, no global information about the network structure was required for signal propagation. RW was used in the previous implementation of discrete-event simulations in Mišić, Sporns, and McIntosh (2014) and Mišić, Goñi, et al. (2014).

In SP, a message or packet was transmitted along the shortest path between the source and destination nodes. If multiple shortest paths existed, a message or packet was randomly transmitted to one of the neighboring nodes closest to the destination node in terms of the shortest path length with equal probability. For signal propagation with this strategy, full connectivity information of the network was necessary to determine the shortest path. Propagation with SP has been assumed in the definitions of several network metrics, e.g., global efficiency (Latora & Marchiori, 2001; Rubinov & Sporns, 2010).

In contrast to SP, iRW only uses local information from neighboring nodes. We implemented three different versions of iRW (Figure 2A): (1) iRW with a rule of avoiding busy nodes (iRW_a_), (2) iRW with a rule of deterministic transmission to the destination node if it is one of the neighboring nodes (iRW_d_), and (3) iRW with both of these rules combined (iRW_a+d_). In iRW_a_ and iRW_a+d_, a message or packet was transmitted to an unoccupied neighboring node if it existed. Otherwise, a message or packet was transmitted to the neighboring node with the shortest queue length in its buffer. If there were multiple such candidate nodes to transmit a message or packet, one of them was randomly selected with an equal probability. In iRW_d_ and iRW_a+d_, a message or packet was deterministically transmitted to the destination node if it was in the set of neighboring nodes to the current node.

In bRW (Avena-Koenigsberger et al., 2019), the propagation behavior changes between that of RW and SP through the control parameter *c* in *p_ijd_* = exp(*−*(*c*(*l_ij_* + *g_jd_*) + *l_ij_*))/*Z_id_*, where *p_ijd_* is the probability that a message or packet was transmitted from the current node *i* to one of its neighboring nodes *j* when *d* is the destination node of the message or packet, *l_ij_* is the length of the edge connecting nodes *i* and *j* (or *l_ij_* = ∞ if this connection is absent), *g_jd_* is the shortest path length between node *j* and destination *d*, and *Z_id_* = ∑*_j_* exp(*−*(*c*(*l_ij_* + *g_jd_*) + *l_ij_*)) is a normalization factor. All edges were assigned the same length of 1 because the structural brain network used in our simulations was a binary network and its distance information was also not available. The term *l_ij_* + *g_jd_* corresponds to the minimum distance in hops between node *i* and destination *d* via neighboring node *j*. The control parameter *c* can change the extent to which the global information about the network structure is available for signal propagation (see Figure 3A for how *c* changes the transition probability in a toy example case). If *c* = 0, the transition probability *p_ijd_* is reduced to that of RW as *p_ijd_* becomes the same across all neighboring nodes. If *c →* ∞, *p_ijd_* is reduced to that of SP as *p_ijd_ →* 1 at a neighboring node *j* for which *l_ij_* + *g_jd_* is the minimum (i.e., *j* closest to *d* in terms of the shortest path length). When there are two or more such neighboring nodes due to the existence of multiple shortest paths, *p_ijd_ →* 1/the number of those nodes.

### Communication metrics

From the simulated signal propagation in the connectome, we computed two nodewise communication metrics: the proportion of the time that a node was busy (*utilization*) and the mean number of messages or packets at a node (*node contents*) as in Mišić, Sporns, and McIntosh (2014) and Mišić, Goñi, et al. (2014). These metrics were derived from the following simulation variables for node *i* at time *t*: the server contents *s_i_*(*t*) *∈ {*0, 1*}*, which represents whether there is a message or packet currently in service (*s_i_*(*t*) = 1) or not (*s_i_*(*t*) = 0), and the queue length *q_i_*(*t*) *∈ {*0, …, *H}* (*H*: the maximum buffer size), which represents the number of messages or packets in the buffer. The utilization of node *i* was defined as the proportion of simulation time having *s_i_*(*t*) = 1. The node contents were defined as the sum of the server and queue contents, *n_i_*(*t*) = *s_i_*(*t*) + *q_i_*(*t*). In the Results section, we show the normalized version of node contents *n_i_*(*t*)/(1 + *H*) to properly compare this metric between message switching and packet switching.

### Surrogate networks

In the surrogate data analysis, we constructed two types of surrogate networks by rewiring edges in the original structural brain network. We considered i) fully randomized surrogate networks in which edges were randomly rewired with no constraint and ii) degree-preserved surrogate networks in which edges were rewired while preserving the sequences of the in-degree and out-degree of nodes in the original brain network. We generated 100 different realizations of the fully randomized or degree-preserved surrogate networks to obtain the distributions of the elapsed time in Figure 5C or D. The same sets of the 100 surrogate networks were used in the simulations of signal propagation for each combination of the propagation strategies and switching architectures.

## Acknowledgements

This work was supported by the Japan Society for the Promotion of Science, KAKENHI Grants JP20H05066 and JP21K15610 to MF.

## Supplementary Results

To confirm our findings in the elapsed time comparisons (Figures 2B and 3B) in the Results section, we examined whether these results were robust against having the following changes applied to the default simulation setting of message switching and packet switching.

i. **No buffer size limit**. The limit on the buffer size in the default setting was removed. Figure S1 shows that the results were essentially the same as those obtained with the buffer size limit.
ii. **Decrease/increase of the number of packets in a message**. The number of packets *n* in a message was changed from five in the default setting to three or ten. While the different values of *n* changed the elapsed time of packet switching, the effects of packetization on the elapsed time (longer under RW and bRW with small control parameter *c*; shorter under iRW and bRW with intermediate *c*; unchanged under SP and bRW with high *c*) remained the same when *c* was decreased or increased (Figures S2 and S3).
iii. **Decrease/increase of the arrival rate**. The arrival rate *λ* was changed from 0.01 in the default setting to 0.005 (less frequent signal generation) or 0.02 (more frequent signal generation), which are the minimum and the maximum arrival rates used in Mišić, Sporns, & McIntosh (2014). As in this previous study, we kept the service rate *µ* unchanged because the system dynamics were dependent on the ratio *λ*/*µ*. Figure S4 shows that the effects of packetization on the elapsed time also held with *λ* = 0.005. When *λ* = 0.02, packetization conversely lengthened the elapsed time under iRW_a_ (Figure S5A). However, the conclusion about the effects of packetization still held since the communication speed of iRW_a_ was much slower than that of the other versions of iRW in this case, and therefore iRW_a_ was no longer a speed-and cost-balanced strategy at *λ* = 0.02. In bRW, while the range of *c* where packetization improved the transmission efficiency became smaller when *λ* = 0.005 (Figure S4B) and larger when *λ* = 0.02 (Figure S5B), the overall trend of the elapsed time difference between message switching and packet switching remained the same.
iv. **No packet overtaking**. The simulation setting was changed to prevent packets from overtaking other packets from the same message. This was realized by first replacing the default assumption, where each packet can individually take its own route, with another one, where all packets in the same message follow the path of their foremost packet. Then, the default last-in-first-out queueing system, in which packets can be overtaken, was replaced with a first-in-first-out system. With this change, the oldest message or packet in a node’s buffer was to start its service first after the service of the previously occupying message or packet at that node was finished, and a new message or packet arriving at a node was removed when its buffer was fully occupied by others. We observed that these changes shortened the elapsed time of packet switching under slow propagation strategies (e.g., RW); however, the effects of packetization on the elapsed time persisted (Figure S6).

## Supplementary Figures

**Figure S1.**
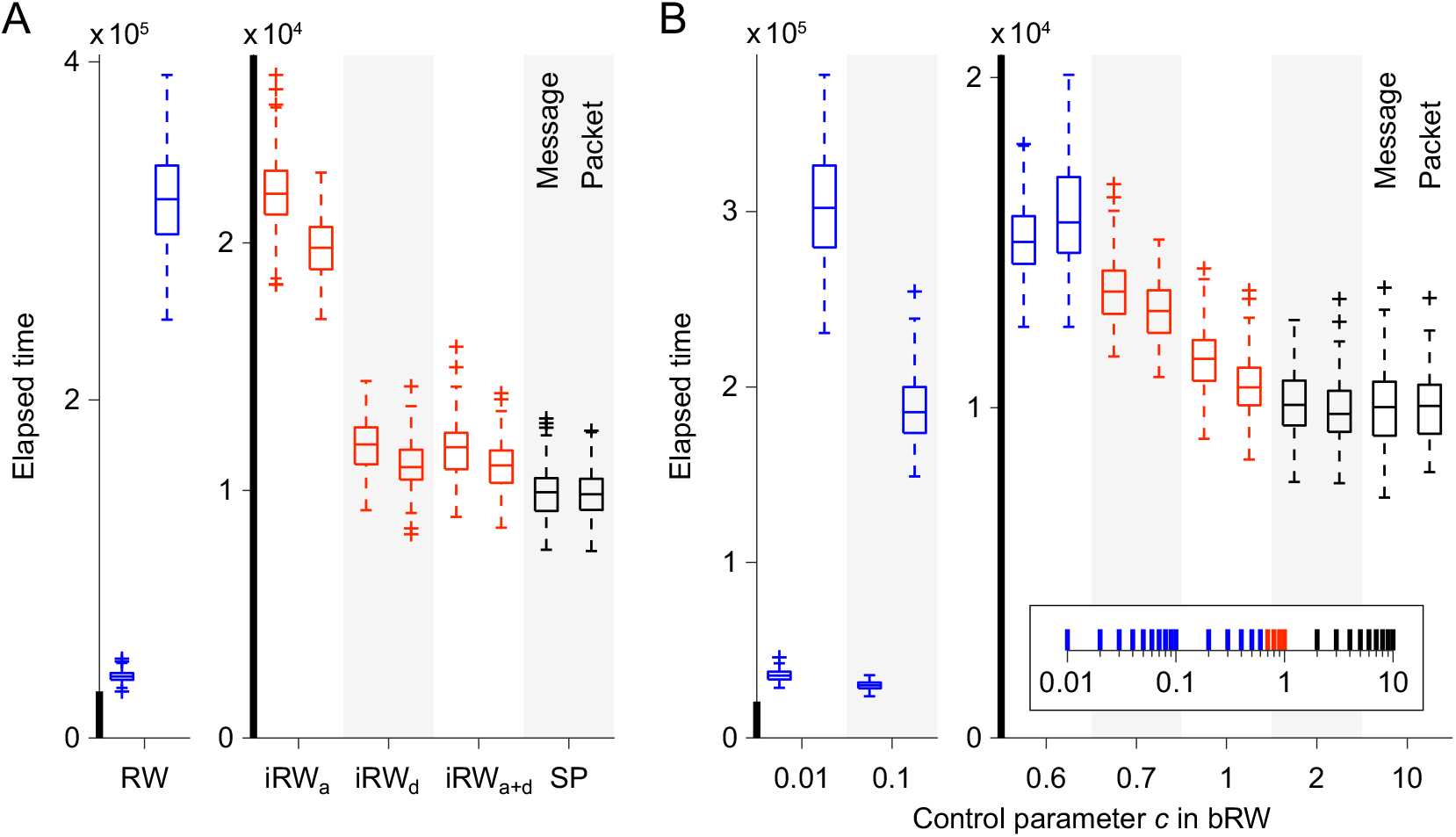
Elapsed time for transmitting 100 messages or packet sets in the simulations with no buffer size limit. (A) Elapsed time with RW, iRW, and SP. (B) Elapsed time with bRW. The boxplots are colored in the same manner as in Figures 2B and 3B.

**Figure S2.**
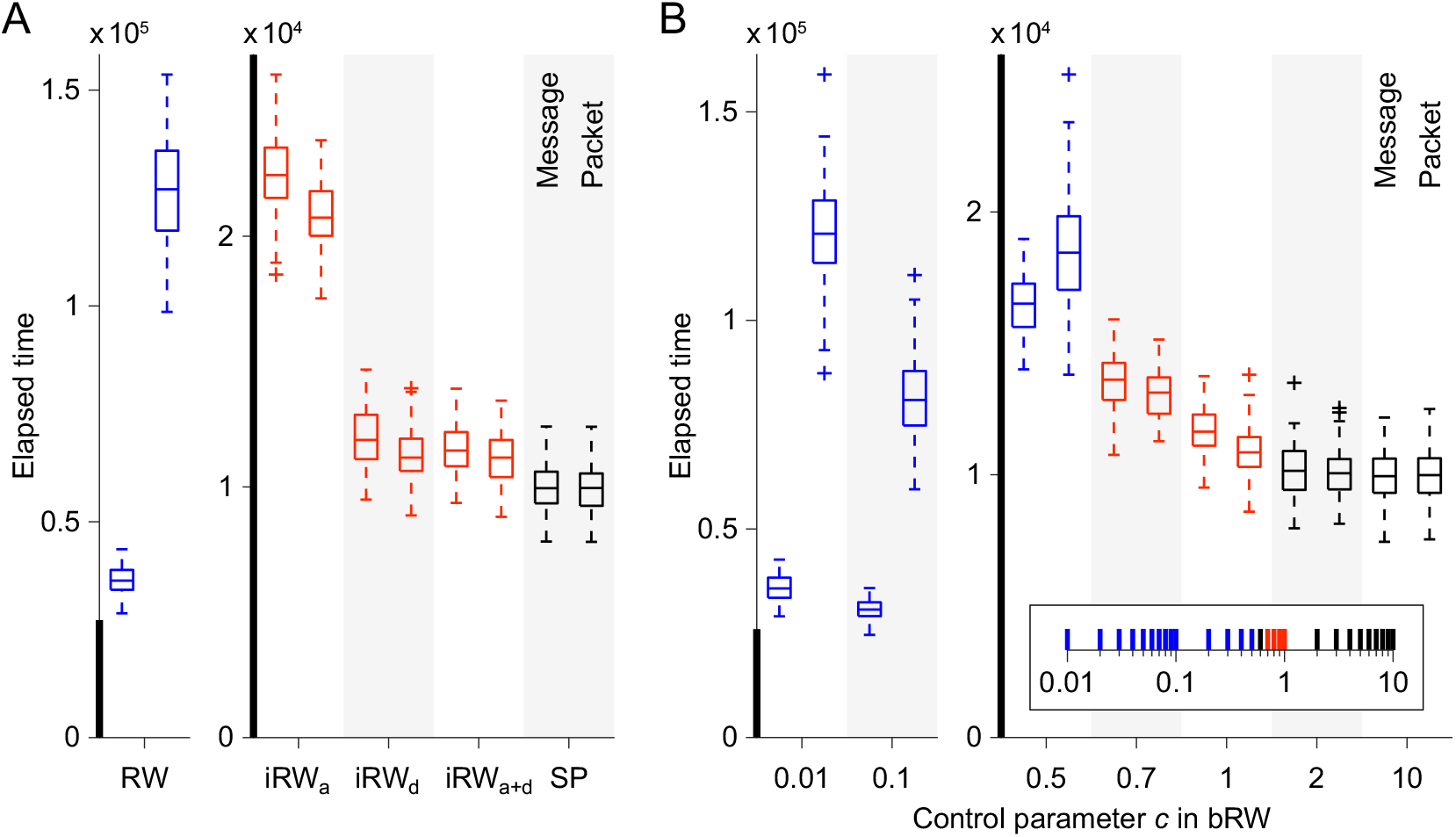
Elapsed time for transmitting 100 messages or packet sets in the simulations with the number of packets in a message *n* = 3. (A) Elapsed time with RW, iRW, and SP. (B) Elapsed time with bRW.

**Figure S3.**
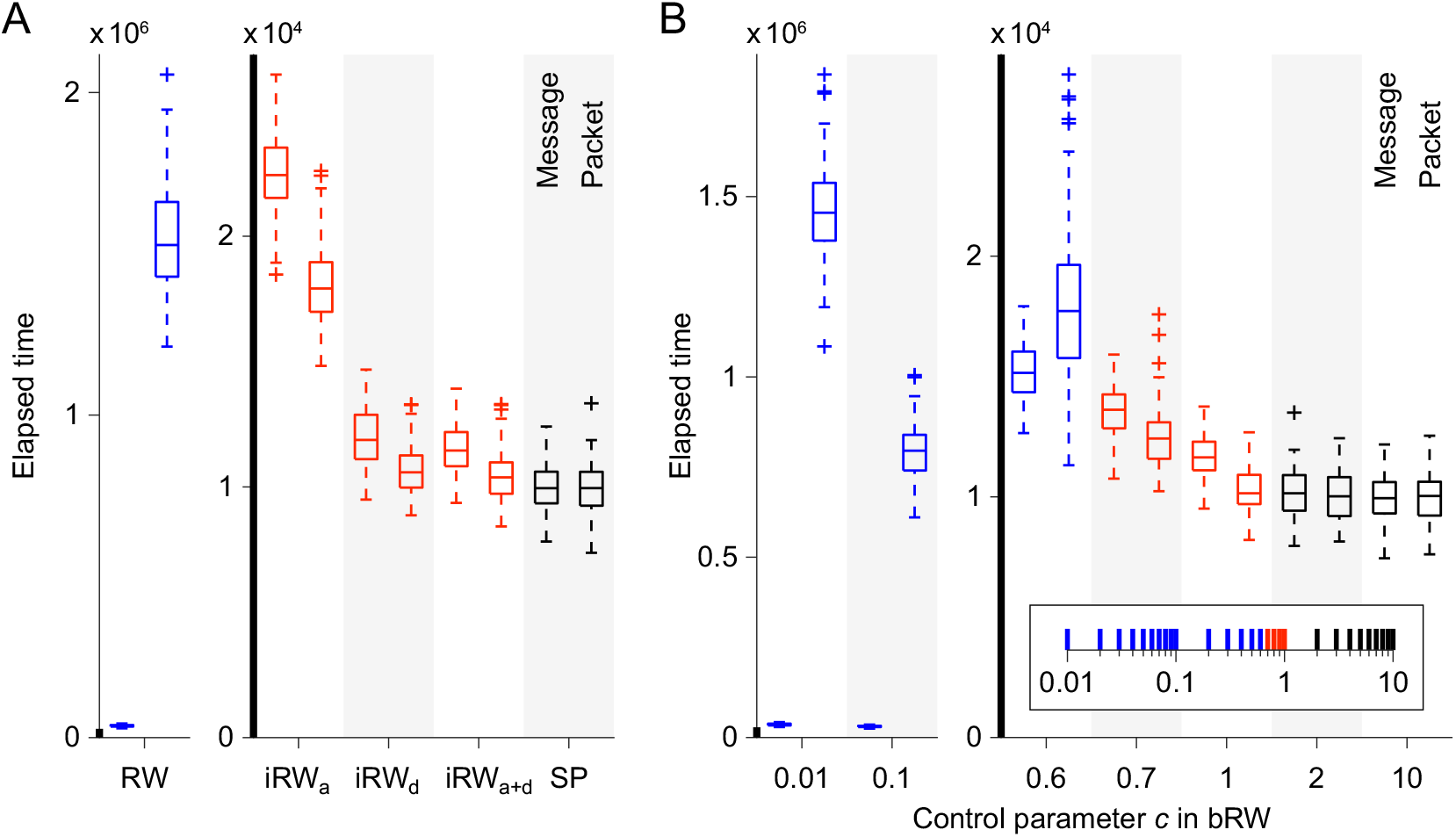
Elapsed time for transmitting 100 messages or packet sets in the simulations with the number of packets in a message *n* = 10. (A) Elapsed time with RW, iRW, and SP. (B) Elapsed time with bRW.

**Figure S4.**
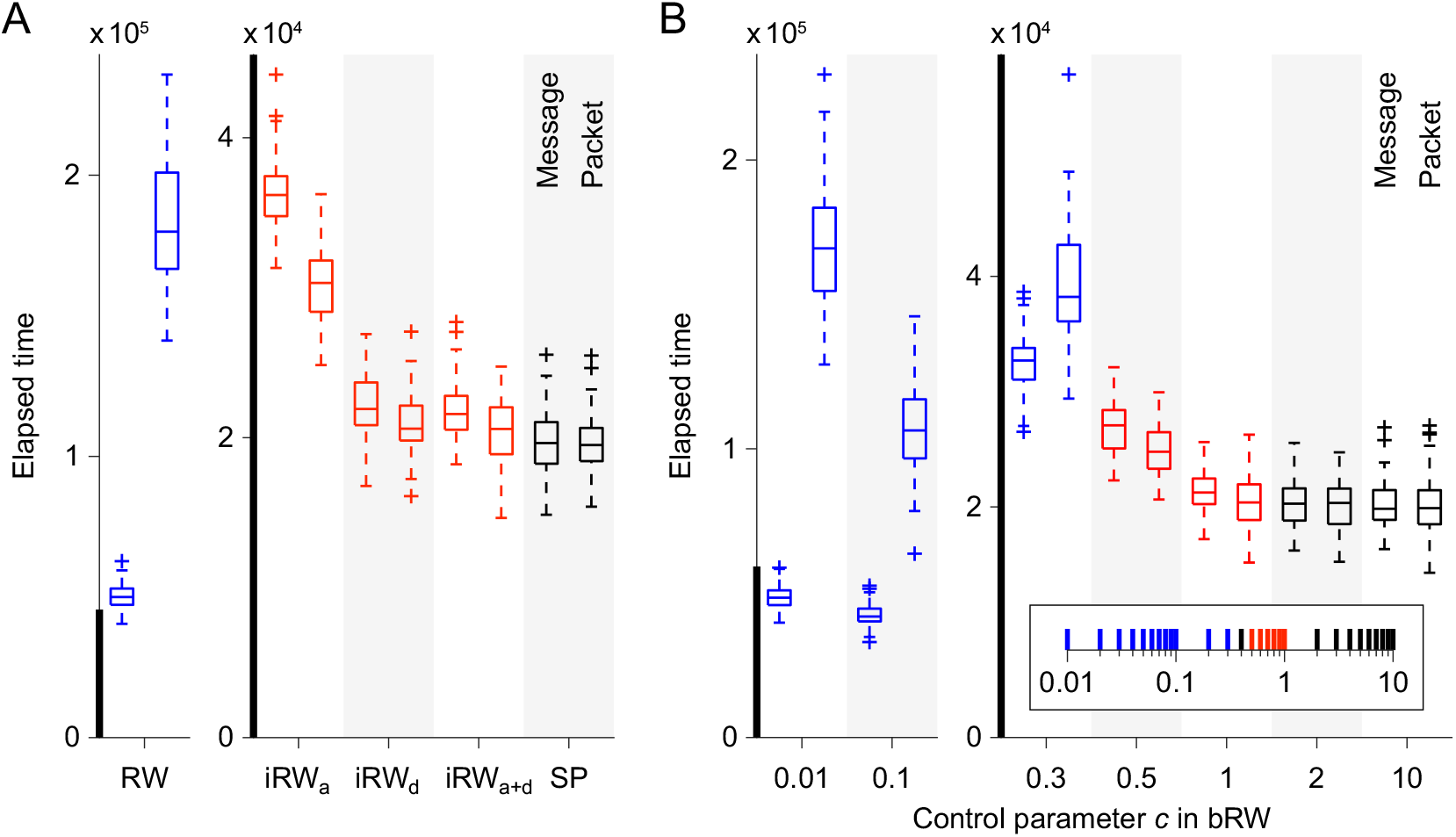
Elapsed time for transmitting 100 messages or packet sets in the simulations with the arrival rate *λ* = 0.005. (A) Elapsed time with RW, iRW, and SP. (B) Elapsed time with bRW.

**Figure S5.**
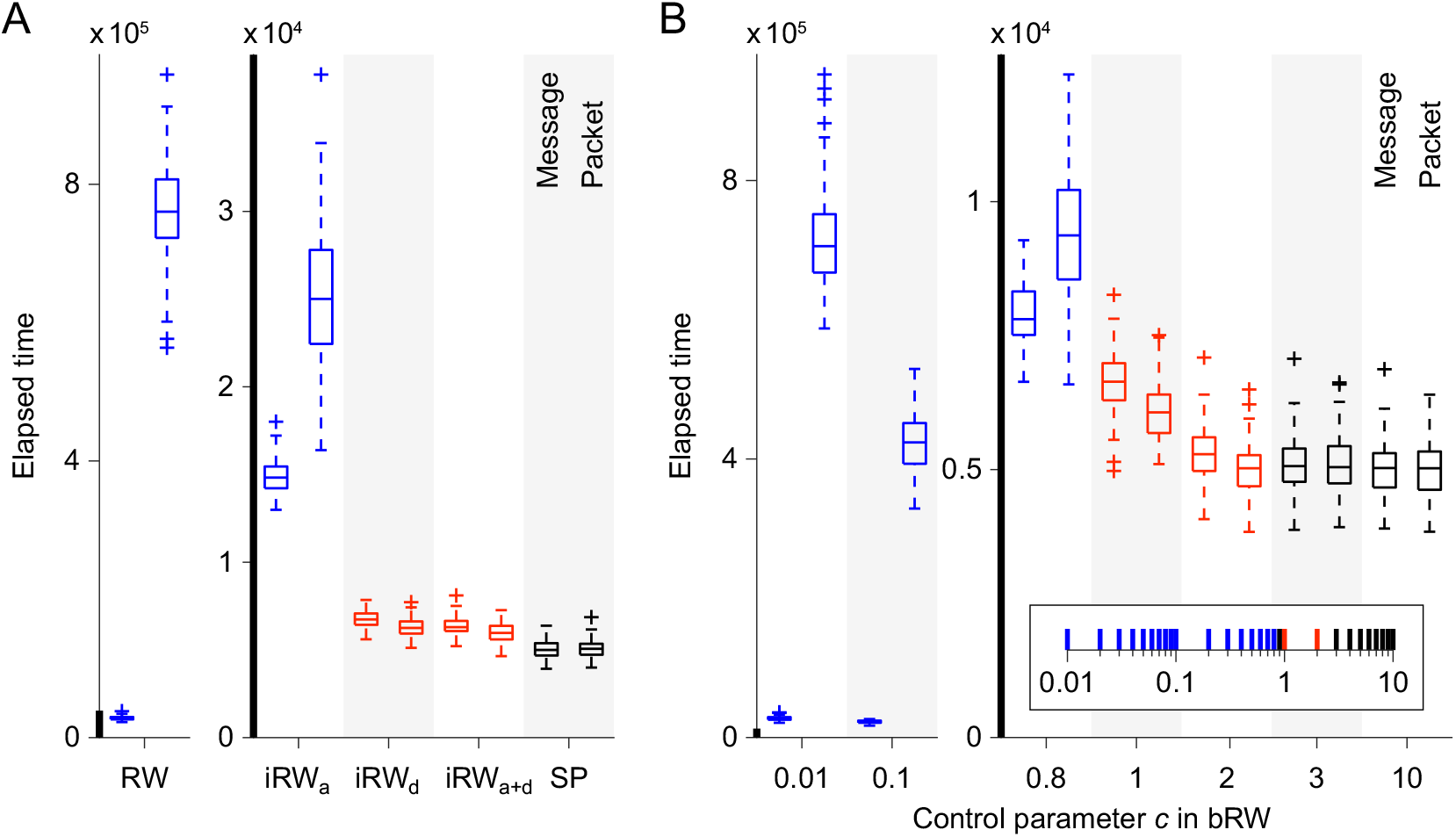
Elapsed time for transmitting 100 messages or packet sets in the simulations with the arrival rate *λ* = 0.02. (A) Elapsed time with RW, iRW, and SP. (B) Elapsed time with bRW.

**Figure S6.**
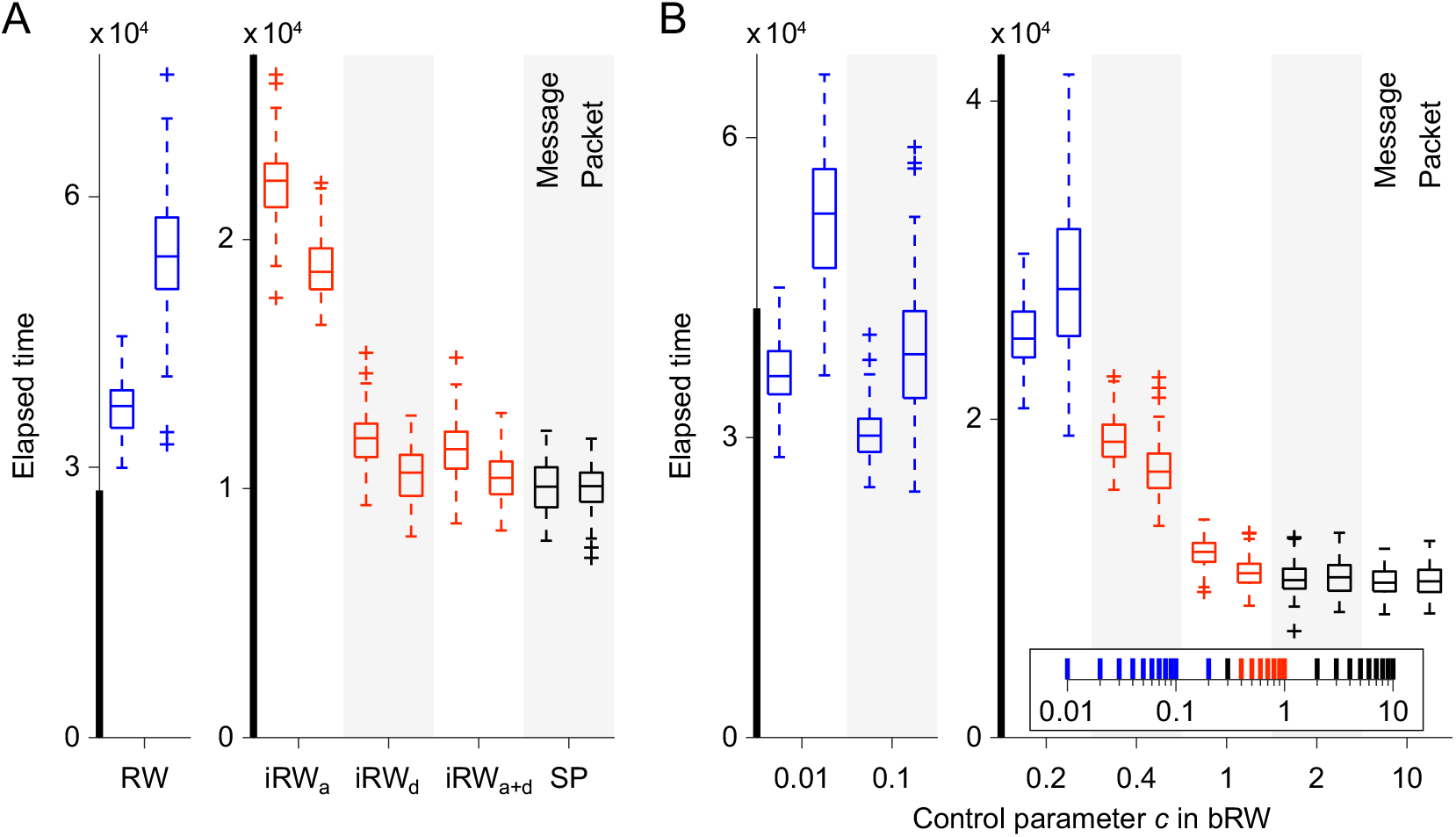
Elapsed time for transmitting 100 messages or packet sets in the simulations with no packet overtaking. (A) Elapsed time with RW, iRW, and SP. (B) Elapsed time with bRW.

**Figure S7.**
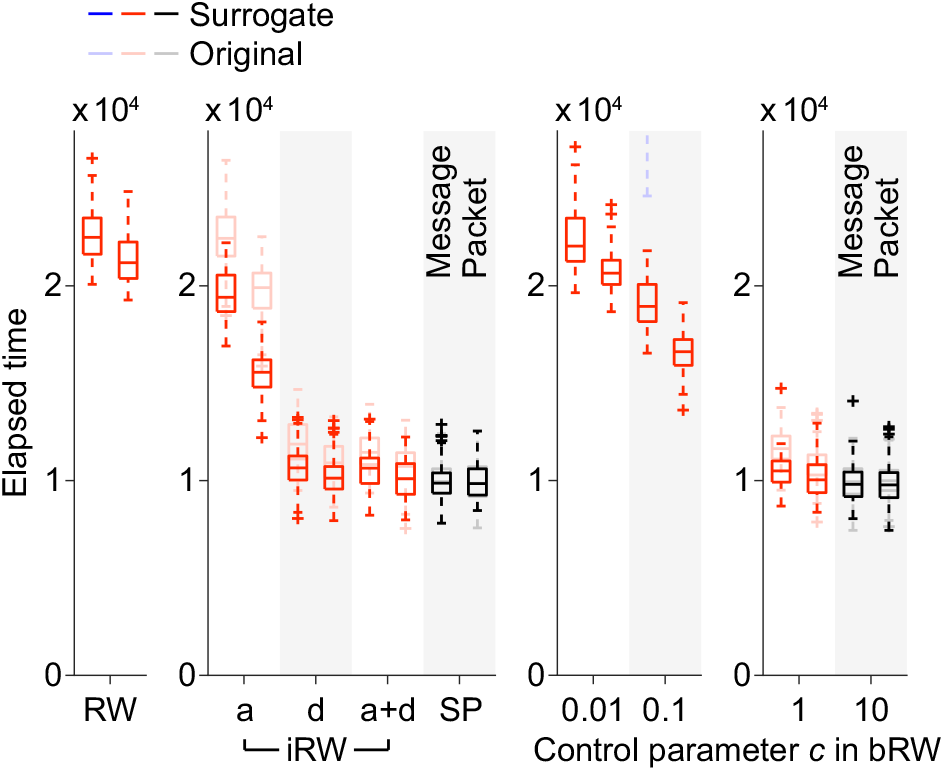
The boxplots zoomed in along the vertical axis for RW and bRW with *c* = 0.1 and 0.01 in Figure 5C. The boxplots for iRW, SP, and bRW with *c* = 1 and 10 are also shown with the same vertical axis range.

## References

Abdelnour, F., Voss, H. U., & Raj, A. (2014). Network diffusion accurately models the relationship between structural and functional brain connectivity networks. NeuroImage, 90, 335–347.

Avena-Koenigsberger, A., Mišić, B., Hawkins, R. X. D., Griffa, A., Hagmann, P., Goñi, J., & Sporns, O. (2017). Path ensembles and a tradeoff between communication efficiency and resilience in the human connectome. Brain Struct. Funct., 222, 603–618.

Avena-Koenigsberger, A., Misic, B., & Sporns, O. (2018). Communication dynamics in complex brain networks. Nat. Rev. Neurosci., 19, 17–33.

Avena-Koenigsberger, A., Yan, X., Kolchinsky, A., van den Heuvel, M. P., Hagmann, P., & Sporns, O. (2019). A spectrum of routing strategies for brain networks. PLOS Comput. Biol., 15, e1006833.

Baddeley, R., Abbott, L. F., Booth, M. C. A., Sengpiel, F., Freeman, T., Wakeman, E. A., & Rolls, E. T. (1997). Responses of neurons in primary and inferior temporal visual cortices to natural scenes. Proc. R. Soc. Lond. B Biol. Sci., 264, 1775–1783.

Banks, J., & Carson II, J. S. (1984). Discrete-event system simulation. New Jersey: Prentice-Hall.

Barlow, H. B. (1956). Retinal noise and absolute threshold. J. Opt. Soc. Am., 46, 634–639.

Bettinardi, R. G., Deco, G., Karlaftis, V. M., Van Hartevelt, T. J., Fernandes, H. M., Kourtzi, Z., . . . Zamora-López, G. (2017). How structure sculpts function: unveiling the contribution of anatomical connectivity to the brain’s spontaneous correlation structure. Chaos, 27, 047409.

Betzel, R. F., Faskowitz, J., Mišić, B., Sporns, O., & Seguin, C. (2022). Multi-policy models of interregional communication in the human connectome. bioRxiv, doi, 10.1101/2022.05.08.490752.

Breakspear, M. (2017). Dynamic models of large-scale brain activity. Nat. Neurosci., 20, 340–352.

Cabral, J., Hugues, E., Sporns, O., & Deco, G. (2011). Role of local network oscillations in resting-state functional connectivity. NeuroImage, 57, 130–139.

Chen, Y., Zhang, Z. K., He, Y., & Zhou, C. (2020). A large-scale high-density weighted structural connectome of the macaque brain acquired by predicting missing links. Cereb. Cortex, 30, 4771–4789.

Crofts, J. J., & Higham, D. J. (2009). A weighted communicability measure applied to complex brain networks. J. R. Soc. Interface, 6, 411–414.

Damoiseaux, J. S., Rombouts, S. A. R. B., Barkhof, F., Scheltens, P., Stam, C. J., Smith, S. M., & Beckmann, C. F. (2006). Consistent resting-state networks across healthy subjects. Proc. Natl. Acad. Sci. U. S. A., 103, 13848–13853.

Deco, G., Cabral, J., Woolrich, M. W., Stevner, A. B. A., van Hartevelt, T. J., & Kringelbach, M. L. (2017). Single or multiple frequency generators in on-going brain activity: a mechanistic whole-brain model of empirical MEG data. NeuroImage, 152, 538–550.

Deco, G., & Jirsa, V. K. (2012). Ongoing cortical activity at rest: criticality, multistability, and ghost attractors. J. Neurosci., 32, 3366–3375.

Faisal, A. A., Selen, L. P. J., & Wolpert, D. M. (2008). Noise in the nervous system. Nat. Rev. Neurosci., 9, 292–303.

Fox, M. D., Snyder, A. Z., Vincent, J. L., Corbetta, M., Van Essen, D. C., & Raichle, M. E. (2005). The human brain is intrinsically organized into dynamic, anticorrelated functional networks. Proc. Natl. Acad. Sci. U. S. A., 102, 9673–9678.

Fukushima, M., & Sporns, O. (2018). Comparison of fluctuations in global network topology of modeled and empirical brain functional connectivity. PLOS Comput. Biol., 14, e1006497.

Fukushima, M., & Sporns, O. (2020). Structural determinants of dynamic fluctuations between segregation and integration on the human connectome. *Commun*. Biol., 3, 606.

Funahashi, S. (2015). Functions of delay-period activity in the prefrontal cortex and mnemonic scotomas revisited. Front. Syst. Neurosci., 9, 2.

Gerraty, R. T., Davidow, J. Y., Foerde, K., Galvan, A., Bassett, D. S., & Shohamy, D. (2018). Dynamic flexibility in striatal-cortical circuits supports reinforcement learning. J. Neurosci., 38, 2442–2453.

Goldman-Rakic, P. S. (1996). Regional and cellular fractionation of working memory. Proc. Natl. Acad. Sci. U. S. A., 93, 13473–13480.

Goñi, J., van den Heuvel, M. P., Avena-Koenigsberger, A., Velez de Mendizabal, N., Betzel, R. F., Griffa, A., . . . Sporns, O. (2014). Resting-brain functional connectivity predicted by analytic measures of network communication. Proc. Natl. Acad. Sci. U. S. A., 111, 833–838.

Graham, D. (2014). Routing in the brain. Front. Comput. Neurosci., 8, 44.

Graham, D. (2021). An internet in your head: a new paradigm for how the brain works. New York: Columbia University Press.

Graham, D., & Rockmore, D. (2011). The packet switching brain. J. Cogn. Neurosci., 23, 267–276.

Hagmann, P., Cammoun, L., Gigandet, X., Meuli, R., Honey, C. J., Wedeen, V. J., & Sporns, O. (2008). Mapping the structural core of human cerebral cortex. PLOS Biol., 6, e159.

Hao, Y., & Graham, D. (2020). Creative destruction: sparse activity emerges on the mammal connectome under a simulated communication strategy with collisions and redundancy. Netw. Neurosci., 4(4), 1055–1071.

Harriger, L., van den Heuvel, M. P., & Sporns, O. (2012). Rich club organization of macaque cerebral cortex and its role in network communication. PLOS ONE, 7, e46497.

Honey, C. J., Kötter, R., Breakspear, M., & Sporns, O. (2007). Network structure of cerebral cortex shapes functional connectivity on multiple time scales. Proc. Natl. Acad. Sci. U. S. A., 104, 10240–10245.

Honey, C. J., Sporns, O., Cammoun, L., Gigandet, X., Thiran, J. P., Meuli, R., & Hagmann, P. (2009). Predicting human resting-state functional connectivity from structural connectivity. Proc. Natl. Acad. Sci. U. S. A., 106, 2035–2040.

Kleinrock, L. (1976). Queueing systems, volume 2: computer applications. New York: Wiley-Interscience.

Kötter, R. (2004). Online retrieval, processing, and visualization of primate connectivity data from the CoCoMac database. Neuroinformatics, 2, 127–144.

Latora, V., & Marchiori, M. (2001). Efficient behavior of small-world networks. Phys. Rev. Lett., 87, 198701.

Liu, Y. (1996). Queueing network modeling of elementary mental processes. Psychol. Rev., 103, 116–136.

Markov, N. T., Ercsey-Ravasz, M. M., Ribeiro Gomes, A. R., Lamy, C., Magrou, L., Vezoli, J., . . . Kennedy, H. (2014). A weighted and directed interareal connectivity matrix for macaque cerebral cortex. Cereb. Cortex, 24, 17–36.

McGill, W. J. (1967). Neural counting mechanisms and energy detection in audition. J. Math. Psychol., 4, 351–376.

Mišić, B., Betzel, R. F., Nematzadeh, A., Goñi, J., Griffa, A., Hagmann, P., . . . Sporns, O. (2015). Cooperative and competitive spreading dynamics on the human connectome. Neuron, 86, 1518–1529.

Mišić, B., Goñi, J., Betzel, R. F., Sporns, O., & McIntosh, A. R. (2014). A network convergence zone in the hippocampus. PLOS Comput. Biol., 10, e1003982.

Mišić, B., Sporns, O., & McIntosh, A. R. (2014). Communication efficiency and congestion of signal traffic in large-scale brain networks. PLOS Comput. Biol., 10, e1003427.

Modha, D. S., & Singh, R. (2010). Network architecture of the long-distance pathways in the macaque brain. Proc. Natl. Acad. Sci. U. S. A., 107, 13485–13490.

Nádasdy, Z., Hirase, H., Czurkó, A., Csicsvari, J., & Buzsáki, G. (1999). Replay and time compression of recurring spike sequences in the hippocampus. J. Neurosci., 19, 9497–9507.

Oh, S. W., Harris, J. A., Ng, L., Winslow, B., Cain, N., Mihalas, S., . . . Zeng, H. (2014). A mesoscale connectome of the mouse brain. Nature, 508, 207–214.

Palmigiano, A., Geisel, T., Wolf, F., & Battaglia, D. (2017). Flexible information routing by transient synchrony. Nat. Neurosci., 20, 1014–1022.

Pope, M., Fukushima, M., Betzel, R. F., & Sporns, O. (2021). Modular origins of high-amplitude cofluctuations in fine-scale functional connectivity dynamics. Proc. Natl. Acad. Sci. U. S. A., 118, e2109380118.

Rubinov, M., & Sporns, O. (2010). Complex network measures of brain connectivity: uses and interpretations. NeuroImage, 52, 1059–1069.

Seguin, C., Tian, Y., & Zalesky, A. (2020). Network communication models improve the behavioral and functional predictive utility of the human structural connectome. Netw. Neurosci., 4, 980–1006.

Seguin, C., van den Heuvel, M. P., & Zalesky, A. (2018). Navigation of brain networks. Proc. Natl. Acad. Sci. U. S. A., 115, 6297–6302.

Sporns, O., Honey, C. J., & Kötter, R. (2007). Identification and classification of hubs in brain networks. PLOS ONE, 2, e1049.

Stephan, K. E., Kamper, L., Bozkurt, A., Burns, G. A. P. C., Young, M. P., & Kötter, R. (2001). Advanced database methodology for the Collation of Connectivity data on the Macaque brain (CoCoMac). Philos. Trans. R. Soc. Lond. B. Biol. Sci., 356, 1159–1186.

Tolhurst, D. J., Smyth, D., & Thompson, I. D. (2009). The sparseness of neuronal responses in ferret primary visual cortex. J. Neurosci., 29, 2355–2370.

Van Essen, D. C., Smith, S. M., Barch, D. M., Behrens, T. E. J., Yacoub, E., & Ugurbil K for the WU-Minn HCP Consortium. (2013). The WU-Minn Human Connectome Project: an overview. NeuroImage, 80, 62–79.

Yeo, B. T. T., Krienen, F. M., Sepulcre, J., Sabuncu, M. R., Lashkari, D., Hollinshead, M., . . . Buckner, R. L. (2011). The organization of the human cerebral cortex estimated by intrinsic functional connectivity. J. Neurophysiol., 106, 1125–1165.

